# End-to-end computational approach to the design of RNA biosensors for miRNA biomarkers of cervical cancer

**DOI:** 10.1101/2021.07.09.451282

**Authors:** Priyannth Ramasami S. Baabu, Shivaramakrishna Srinivasan, Swetha Nagarajan, Sangeetha Muthamilselvan, Raghavv R. Suresh, Thamarai Selvi, Ashok Palaniappan

**Affiliations:** Department of Bioengineering, School of Chemical and Biotechnology, SASTRA Deemed University, Thanjavur, Tamil Nadu - 613 401. India; Department of Bioinformatics, School of Chemical and Biotechnology, SASTRA Deemed University, Thanjavur, Tamil Nadu - 613 401. India

**Author notes:** These authors contributed equally (Priyannth Ramasami S. Baabu, Shivaramakrishna Srinivasan). School of Advanced Materials Science and Engineering, Sungkyunkwan University (SKKU), Suwon, 16419, Republic of Korea. Life Sciences Division, TCS Research, Tata Consultancy Services, Chennai, Tamil Nadu 600042. India.

**Keywords:** Cervical cancer biosensor, miRNA biomarker, synthetic toehold switches, genetic circuit design, reaction network modelling, toehold efficacy modelling, toehold switch grammar, machine learning

## Abstract

Cervical cancer is a global public health subject as it affects women in the reproductive ages, and accounts for the second largest burden among cancer patients worldwide with an unforgiving 50% mortality rate. Poor awareness and access to effective diagnosis have led to this enormous disease burden, calling for point-of-care, minimally invasive diagnosis methods. Here, an end-to-end quantitative approach for a new kind of diagnosis has been developed, comprising identification of optimal biomarkers, design of the sensor, and simulation of the diagnostic circuit. Using miRNA expression data in the public domain, we identified circulating miRNA biomarkers specific to cervical cancer using multi-tier screening. Synthetic riboregulators called toehold switches specific for the biomarker panel were then designed. To predict the dynamic range of toehold switches for use in genetic circuits as biosensors, we developed a generic grammar of these switches, and built a multivariate linear regression model using thermodynamic features derived from RNA secondary structure and interaction. The model yielded predictions of toehold efficacy with an adjusted R^2^ = 0.59. Reaction kinetics modelling was performed to predict the sensitivity of the second-generation toehold switches to the miRNA biomarkers. Simulations showed a linear response between 10nM and 100nM before saturation. Our study demonstrates an end-to-end workflow for the efficient design of genetic circuits geared towards the effective detection of unique genomic signatures that would be increasingly important in today’s world. The approach has the potential to direct experimental efforts and minimise costs. All resources including the machine learning toolkit, reaction kinetics simulation, designed toehold sequences, genetic circuits, data and sbml files for replicating and utilizing our study are provided open-source with the iGEM Foundation (https://github.com/igem2019) under GNU GPLv3 licence.

## 1. Introduction

Cervical cancer is the second most common cancer affecting women worldwide [1–3], with 20% of cases in India [4,5]. Most cervical cancers could be attributed to the HPV16 and HPV18 strains of Human Papilloma Virus (HPV) [1,2]. Lifestyle factors also contribute to the etiology of the disease [2]. Cervical cancer tumorigenesis begins with the metaplastic epithelium being infected by HPV strains at the cervical transitional zone, followed by the persistence of the virus, then the progression of the epithelium cells to pre-cancer stage, and finally the progression to cancer [6]. Pap smear test has been a widely used diagnostic method but an invasive, expensive, and time-consuming one [4,7]. Alternative testing strategies are necessary [8–10], and micro-RNAs (miRNAs) circulating in the blood have emerged as effective biomarkers in many cases [11–14]. Coupling data-driven methods for the identification of reliable early-stage diagnostic / prognostic biomarkers with the design of biosensor offers potential for the development of a new generation of molecular diagnostics [15–18].

Synthetic biology is delivering on its promise of harnessing nature’s inherent diversity for the betterment of human health [19]. Bioelectronic diagnostic devices comprise an integrated single-unit reaction chamber housing the sensor element with the necessary reagents for analyte detection, and the corresponding transduction module for providing a reaction readout with optical, piezoelectric, or electrochemical means [20–22]. Toehold switches are a class of synthetic riboregulators with excellent biosensor properties, namely specificity, modularity and orthogonality [23], and such sensors that could sense virtually any RNA sequence [23]. could be freeze-dried on a microfluidic platform for biomarker detection using a cell-free colorimetric assay [24]. Toehold-switch riboregulators are mRNA elements which function by occluding the ribosome from translating a downstream gene (Fig. 1(a)). Upon base pairing with the trigger RNA sequence, the toehold structure of the switch unfolds, permitting expression of a reporter gene. The toehold-switch concept was extensively utilised in the design of a biosensor for detecting Zika virus infection [25,26]. Takahashi *et al.* designed a toehold switch sensor that could detect the conserved region of *C. difficile* in a paper based platform [27]. In this work, we have formulated an alternative testing strategy for cervical cancer based on data-driven identification of biomarker miRNAs, design of cognate toehold-based sensors, and predictive modelling of circuit performance. This synthetic biology approach is the first integrated pathway connecting data from biomarkers to biosensor design and modelling. Software associated with the study is hosted at https://github.com/igem2019, and Supplementary Data could be found at https://doi.org/10.6084/m9.figshare.14915619.v1.

**Fig 1:**
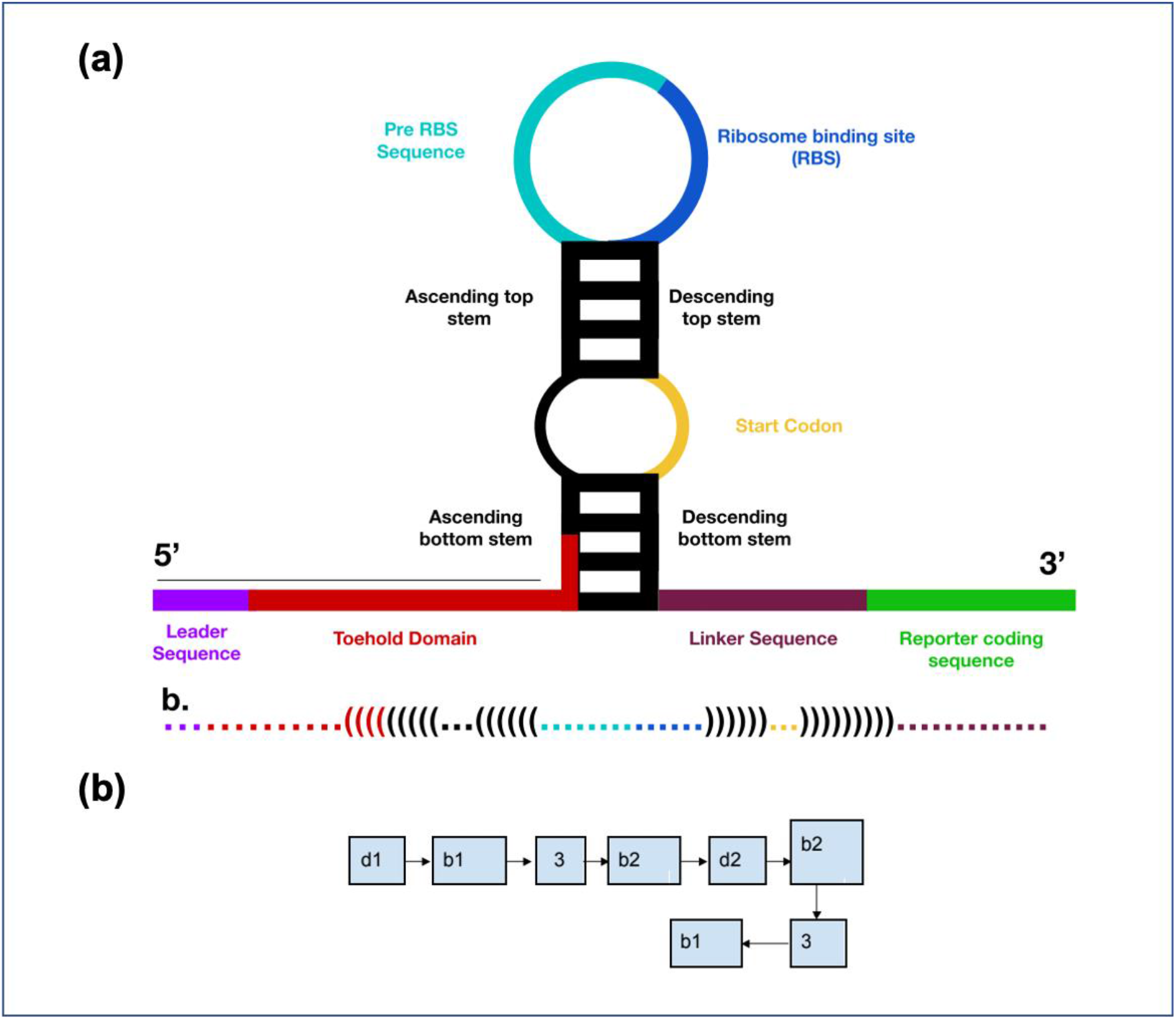
(a) Anatomy of a toehold switch (b) Generic grammar of toehold switch structure for domain parsing, in dot bracket notation.

## 2. Methods

The overall strategy is outlined in Table 1, and discussed in detail below.

**Table 1.** Iterative Design, Build, Test & Learn (DBTL) cycles deployed for developing toehold-switch biosensors specific for a panel of miRNA biomarkers of early-stage cervical cancer.

| **DBTL phase** | Iteration 0:<br>**Biomarker signature development** | Iteration 1:<br>**Optimal toehold switch design** | Iteration 2:<br>**Reaction network modelling of circuit performance** |
| --- | --- | --- | --- |
| **DESIGN** | TCGA data-driven differential expression analysis, survival analysis | Machine learning to design optimal toeholds for identified biomarkers | Modelling the designed circuit using chemical and biochemical kinetics |
| **BUILD** | Cancer vs. control linear models, univariate K-M plots, multivariate Cox regression models (using R) | Feature engineering of toehold datasets using ViennaRNA [28] and in-house Python and shell scripts [29] | Generalising the model to second-generation toeholds with anti-miRs, to inform the design of experiments |
| **TEST** | Models evaluated using p-values and log fold-change of differential expression, and p-values of prognosis models. | Multivariate regression model of toehold efficacy evaluated using $R^2$ goodness of fit (using R) | Validation of predictive modelling |
| **LEARN** | Filtering to four biomarkers, then to three, and finally to two | Optimised toehold switch sequences | Sensitivity, onset of saturation, and specificity of each sensor |

### 2.1 Biomarker Panel Development

The Cancer Genome Atlas (TCGA) provided the miRNA-Seq dataset for normal tissue and cervical adenocarcinoma samples [30]. The dataset contained 312 samples of which 309 were tumour samples and 3 were control samples (tumour-normal matched). The associated clinical data annotated the stage information in the attribute ‘patient.stage_event.clinical_stage’. The number of samples in stages I, II, III and IV were found to be 163, 70, 46 and 21 respectively. The patient sample barcode encoded in variable *hybridisation_REF* was parsed, and the samples were correspondingly annotated as cancer and normal.

RSEM-normalized Illumina HiSeq miRNASeq gene expression data [31] was log_2_ transformed, and analyzed for differential expression using R (https://www.r-project.org/) *limma* package [32]. Linear modelling of stage-annotated gene expression matrix was performed, followed by empirical Bayes adjustment to obtain moderated t-statistics. To account for multiple hypothesis testing and the false discovery rate, the p values of the F-statistic of linear fit were adjusted using the BH method [33]. Differentially expressed miRNAs were considered significant if the absolute log-fold changes > 1.5x and the p-value was <0.05 [34]. A contrast analysis for identifying stage-specific DE miRNAs was then carried out. Finally, the association between the significant early-stage DE miRNAs and the overall survival of a patient was evaluated by univariate Cox proportional hazards regression analysis, and the significant prognostic miRNA biomarkers were retained. All analysis was done R *survival* package [35].

### 2.2 Toehold Switch Design

#### 2.2.1 Toehold switch grammar

A toehold structure consists of two hairpin loops and two base paired stems, that together serve as translation regulator by occluding the RBS in the absence of the cognate trigger molecule (‘ground state’) [23] (Fig. 1). Upon binding of the cognate miRNA (i.e, the trigger sequence) to the toehold domain, the bottom stem unravels and the double-looped structure of the toehold switch collapses to a linear structure, which exposes the sequestered start codon to expression of the reporter gene. Each toehold switch is a modular entity with distinct sequence domains that could be engineered *de novo* for a particular trigger RNA. Toehold switches hold great potential as sensors for exact sequences of RNA, and are custom designed to hybridise with the cognate molecules. Table 2 describes the regions from 5’ to 3’ in the generic toehold switch grammar illustrated in Fig. 1(b).

**Table 2.** Dot-bracket notation of the generalised toehold switch grammar.

| Region | Description |
| --- | --- |
| d1 dots | Toehold domain - not part of bottom stem |
| b1 open brackets | Toehold domain - part of bottom stem, based paired with linker |
| 3 dots (first occurrence) | Sequestered nucleotides opposite to start codon, forming secondary loop |
| b2 open brackets | Ascending top stem |
| d2 dots | Primary loop containing pre-RBS and RBS |
| b2 closed brackets | Descending top stem |
| 3 dots (second occurrence) | Start codon |
| b1 closed brackets | Linker sequence base paired with toehold domain. This leads to reporter gene of interest |

The following preliminary criteria were used in the rational design of toehold sequences:

1. To allow effective binding of the trigger to the switch, the horizontal portion of the toehold domain was longer than 10 nucleotides.
2. The descending bottom stem of the toehold switch was fixed at 12 nucleotides. maintaining switch stability in the OFF state as well as the ORF after the start codon.
3. The canonical RBS sequence AGAGGA was used.
4. The primary loop was made adenine-rich to maintain a larger loop structure.
5. The horizontal linker after the bottom stem was retained as AACCUGGCGGCAGCGCAAAAG, the 21nt long linker used in Green et al. [23].
6. Mature miRNAs are about 22 nts, and might include a stop-codon trinucleotide towards the 5’ region of their sequence. The presence of such stop-codon subsequences could halt the expression of the downstream GFP. Second-generation toehold switches are one solution to this problem [36]. In addition to binding the miRNA, the toehold switch is required to bind another molecule, the anti-miRNA (antimiR) at its 3’ end. AntimiRs are antisense RNA that are complementary and bind to 12 nucleotides at the 3’ end of the toehold switch. Essentially, the trigger is represented by a hybrid molecule with a base-paired region and two free ends. The design of the antimiR sequence could be used in improving the trigger binding event.

#### 2.2.2 Predictive modelling of toehold switch efficacy

The dynamic range describes the sensitivity of the toehold switch, and is the ratio between the maximum and minimum measurable intensities of reporter protein (typically GFP) expression, also known as ON/OFF ratio [23]. The dynamic range of a toehold designed for exact sequences of trigger RNA is the key indicator of its effectiveness, and is generally unknown for new toehold designs. The problem can be addressed using supervised machine learning on toehold-sequence datasets with available dynamic range responses. A previous model developed by CUHK iGEM 2017 offered a web tool for predicting the dynamic range of toehold switches with a modest goodness of fit (R^2^ ~0.2) [37]. To improve on this, we constructed a multivariate linear regression model using the generic grammar of the toehold switch and features engineered from sequence and secondary structure. A validated dataset of the toehold switches with the dynamic range response was available from Green et al. (2014) [23] and Pardee et al (2014, 2016) [24,25]. A total of 230 toehold switch sequences with ON/OFF ratios were thus obtained. The toehold sequence of each instance was parsed into its sub-domains using the generic grammar presented in Fig 1(b). The dot-bracket notation is a representative grammar RNA secondary structure using dots ‘.’ to represent unpaired bases, and matching parentheses ‘(’ and ‘)’ to represent paired bases..additional considerations. 228 toehold instances that conformed to the grammar in Fig 1B were parsed into dot-bracket sequence regions using an in-house regex script (available at https://github.com/igem2019) [29].

Some considerations in domain parsing included:

1. Wobble base pair between G-U in the secondary loop containing the start codon. Wobble base pairs offered better stability in the OFF state and these forward engineered designs could be less leaky
2. Base-paired toehold domains,
3. Base-paired linker regions.
4. Possible unpaired bases in the top stem.

The sequence and structure features used are presented in Table 3. All structural features were calculated using the ViennaRNA suite.

**Table 3.**
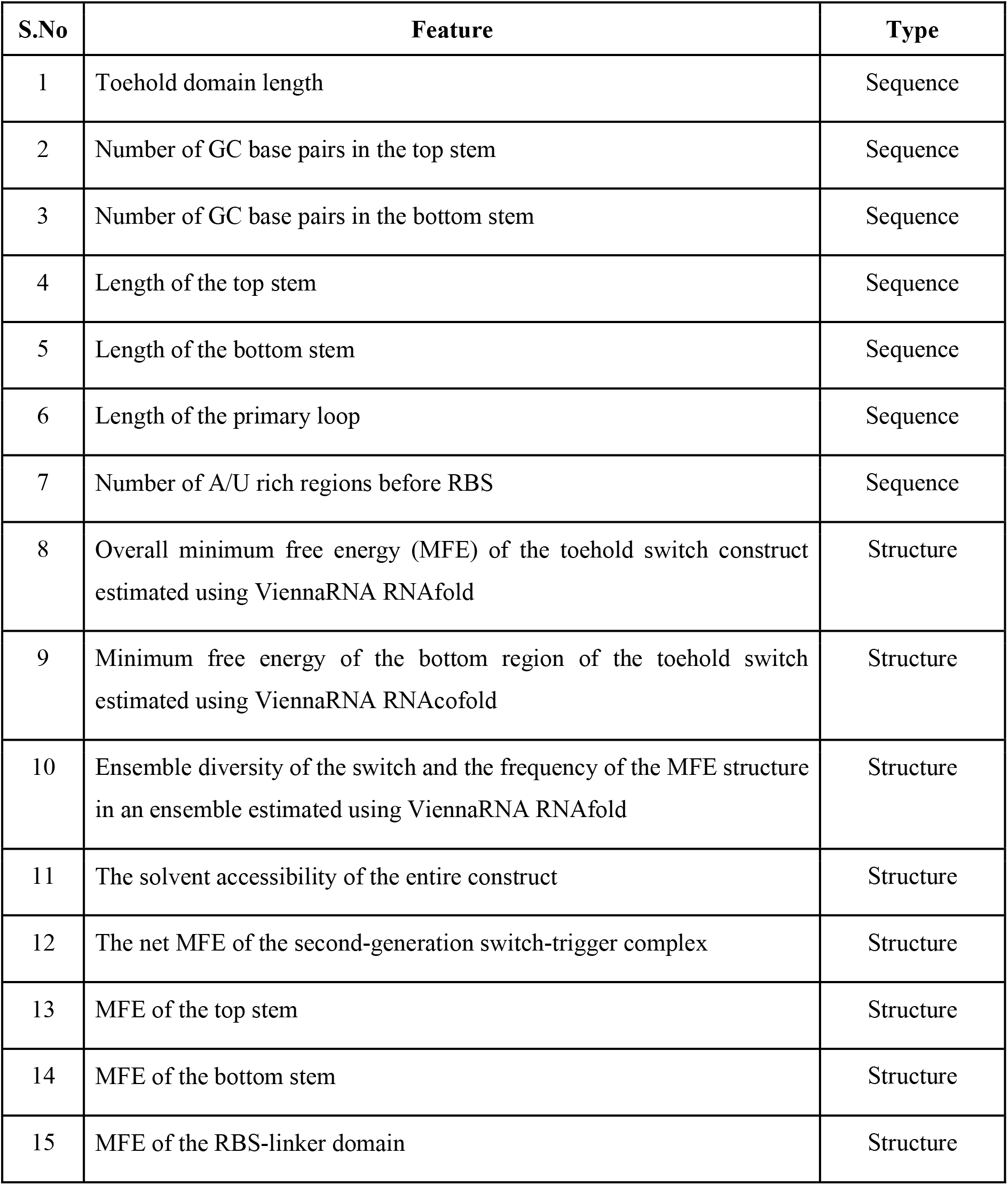
Sequence and structural features of interest in the predictive modelling of a toehold switch.

Feature selection was performed by removing features correlated with Pearson’s ρ > 0.85 were removed. Following a 80:20 train:test split, a multivariate linear regression model was built with the selected features on the train data, and evaluated on the test data for significance and goodness-of-fit, using Python scikit-learn (https://scikit-learn.org/). The scripts and machine learning model are available in the Supplementary Information [29].

#### 2.2.3 gBlock Design

gBlocks are medium-sized genetic constructs capable of housing one or more genes [38], and deposited in the Registry of Biological Parts [39]. Three independent gBlocks were designed using the standard BioBricks (https://biobricks.org), including the constitutive *E. coli* T7 promoter, a 5’ prefix sequence with EcoRI and XbaI recognition sites, and a 3’ suffix sequence with SpeI and PstI recognition sites (Fig. 2). The reporter gene used was GFP-mut3b, capable of enhanced fluorescence due to three point mutations relative to wild GFP. We found that splitting our parts over multiple gBlocks offered less error-prone and faster synthesis. The gBlocks are available in the Supplementary Material.

**Fig 2:**
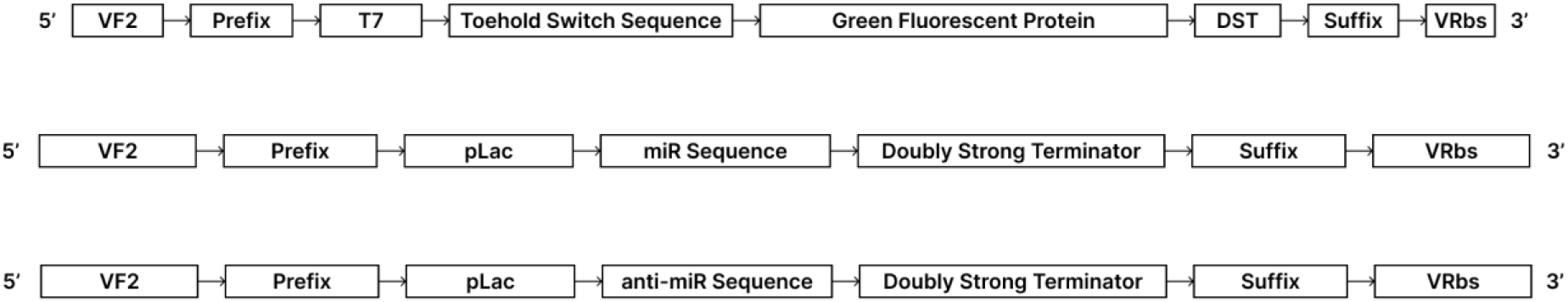
Three gBlocks were designed. (i) Toehold switch-GFP construct; (ii) expression system for miR oligo; and (iii) expression system for antimiR oligo. To allow amplification of the gBlocks, the 5’ prefix is concatenated with the VF2 forward primer, and the 3’ suffix is concatenated with the VR reverse primer. pLac promoters are specifically inducible with lactose.

### 2.3 Reaction network modelling

The chemical and biochemical kinetics model was developed by adding the following features to the first generation toehold switch model developed by CLSB-UK iGEM 2017 [40]:

i. interaction of transcribed miR and antimiR oligos to form the miR-antimiR complex with two free ends, which then binds with the toehold switch
ii. maturation kinetics of the expressed GFP. This generalized model can be presented as:

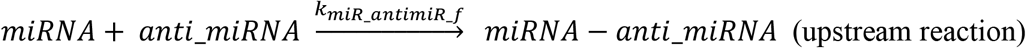

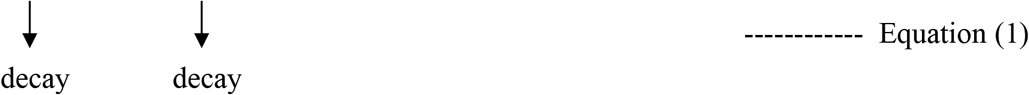

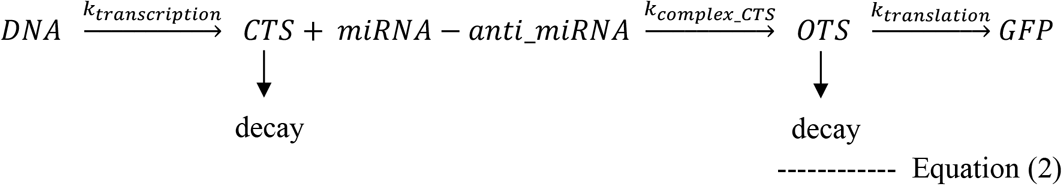

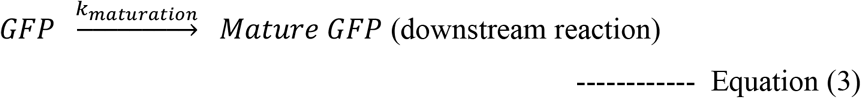

The corresponding ODEs can be given by:

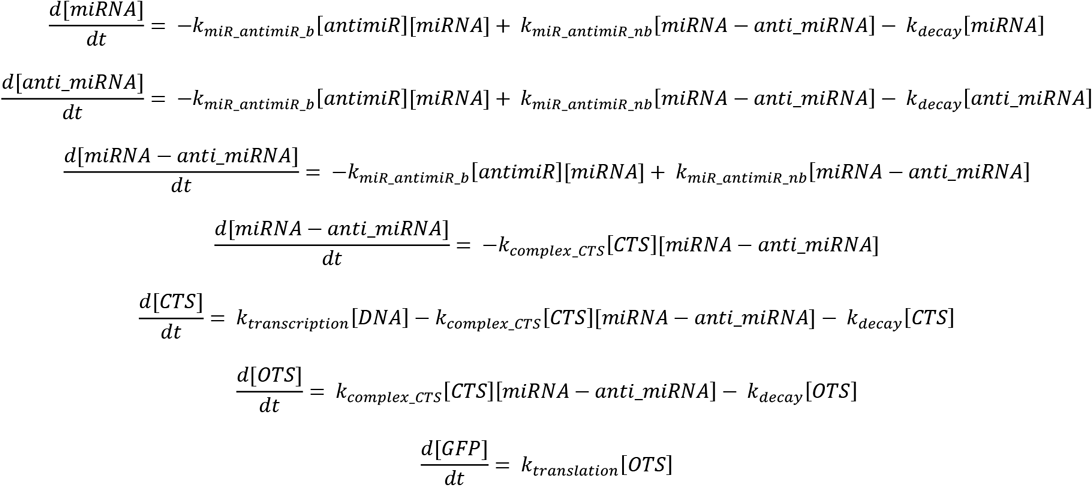

The kinetic parameters for the transcription, translation and maturation reactions in the genetic circuit were identified from literature [40,41]. The kinetics for the miR-antimiR complex formation were modelled assuming a steady state with negligible backward flux, so that the equilibrium constant approximates the forward rate constant at equilibrium. The equilibrium constant for the complex formation is defined as:

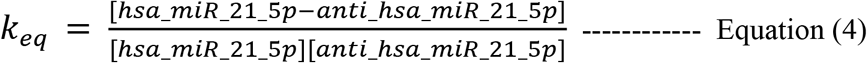

NUPACK was used to obtain the equilibrium concentration values of the miR, antimiR and their complex [42]. The chemical and biochemical kinetics were simulated using MATLAB R2017b (https://www.mathworks.com/).

## 3. Results

### 3.1 Omics Analysis

The top 100 miRNAs of the linear model were screened for log-fold-change with respect to the control samples, stage-specificity, and significance of the trend, and finally corroborated with the literature evidence. This analysis yielded four DE miRNAs (Table 4) [12,43]. The full results of the differential expression analysis and linear modelling are presented in Files S1 and S2 respectively. It is seen that has-miR-21-5p is the most significantly differentially expressed miRNA in squamous cell cervical cancer. It is interesting to note that all four miRNAs are upregulated, highlighting their role as oncomiRs (File S3).

**Table 4.** Putative DE miRNA biomarkers of cervical cancer. LogFC (log-fold change) and adj. p-values with respect to the control as estimated by the linear model are shown, along with the stage-specificity (log-fold change and significance of contrast analysis) of the respective miRNA.

| DE miRNAs | logFC | Adj. p value | Stage_1 - Control | Stage_2 – Control | Stage_3 – Control | Stage_4 – Control | Stage-specificity |
| --- | --- | --- | --- | --- | --- | --- | --- |
| hsa-miR-21-5p | 15.75 | 3E-151 | 2.63 | 2.63 | 2.75 | 2.65 | 4E-08 |
| hsa-miR-20a-5p | 6.60 | 5E-26 | 2.01 | 2.02 | 2.22 | 2.20 | 1E-02 |
| hsa-miR-29a-3p | 13.06 | 1E-70 | 1.99 | 2.31 | 2.18 | 2.64 | 6E-03 |
| hsa-miR-200a-3p | 3.27 | 9E-07 | 6.60 | 6.43 | 6.32 | 6.63 | 3E-17 |

### 3.2 Survival Analysis

The differential expression profiling of miRNAs associated with early-stage cervical cancer identified potential biomarkers, which were further investigated using survival analysis with the Cox proportional hazards model and Kaplan-Meier curves. The biomarkers were subjected to a univariate survival analysis and the model significance was estimated using the log rank test [28]. Fig. 3 shows the K-M plots of the univariate survival analysis for the 4 differentially expressed miRNAs. It could be seen that hsa-miR-29a-3p was not significant at 95% (p-value > 0.05), and was consequently dropped from the hazard ratio estimation (Table 5).

**Table 5.** Hazard ratio (HR), Z-score and p value for the chosen miRNA biomarkers. A HR > 1 indicates a covariate that is positively associated with the event probability (negatively with survival).

| Gene Symbol | HR | Z-score | p-value |
| --- | --- | --- | --- |
| hsa-miR-20a-5p | 0.523 | -2.479 | 0.013 |
| hsa-miR-200a-5p | 1.808 | 2.076 | 0.038 |
| hsa-miR-21-5p | 0.595 | -1.961 | 0.050 |

**Fig 3.**
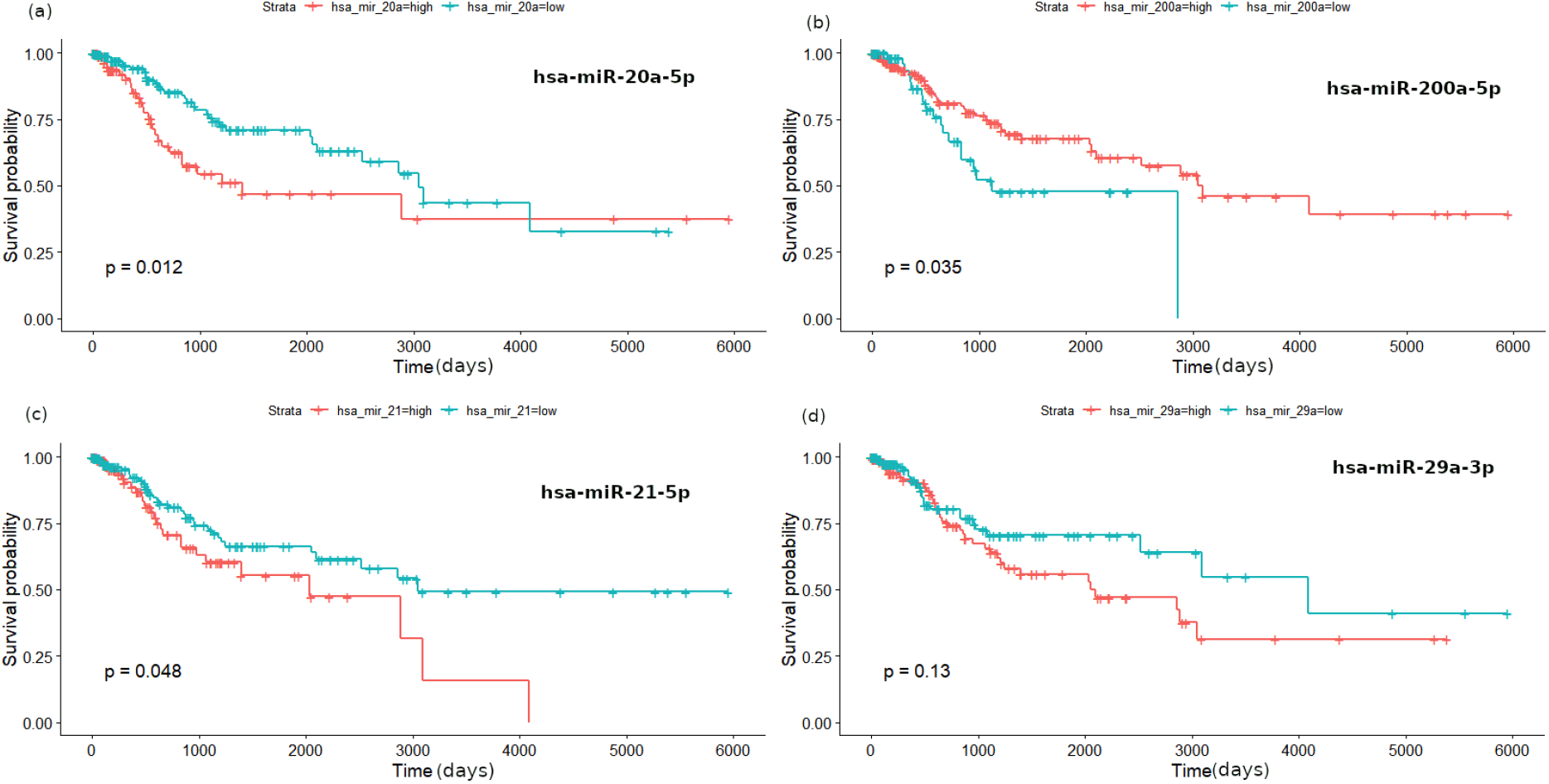
K-M plot for (a) hsa-miR-20a-5p, (b) hsa-miR-200a-5p, (c) hsa-miR-21-5p and (d) hsa-miR-29a-3p.

Next, we constructed a survival risk-score using the multivariate Cox regression model [21], and obtained eqn (5).

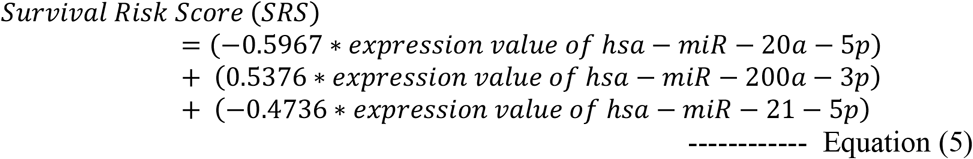

The coefficients of hsa-miR-20a-5p and hsa-miR-21-5p are negative, indicating a protective effect of these miRNAs. Finally, a biomarker panel of hsa-miR-20a-5p, hsa-miR-21-5p and hsa-miR-200a-3p, where the 307 patients who were considered for our analysis was used to classify patients into high-risk and low-risk groups using an optimal cut point identified with the `maxstat` statistic of *survminer* R package. Overall survival curves were generated using the K-M method, and two-sided log rank tests were used to compare the differences in overall survival between the two risk groups. The K-M curve shown in Fig. 4 (a), indicated that the biomarker panel with the three miRNAs was significant (p-value < 0.001). We also evaluated a biomarker panel of just the two miRNAs with protective effect, hsa-miR-20a-5p and hsa-miR-21-5p, considering the significance of hsa-miR-21-5p holds in literature [12,43–48]. The K-M curve shown in Fig. 4 (b), indicated that the significance of this panel was as good as the panel with three miRNAs (p-value < 0.001). From all the above considerations, we settled on the two-marker panel, of hsa-miR-20a-5p and hsa-miR-21-5p, for the prognostic classification of early-stage cervical cancer.

**Fig 4.**
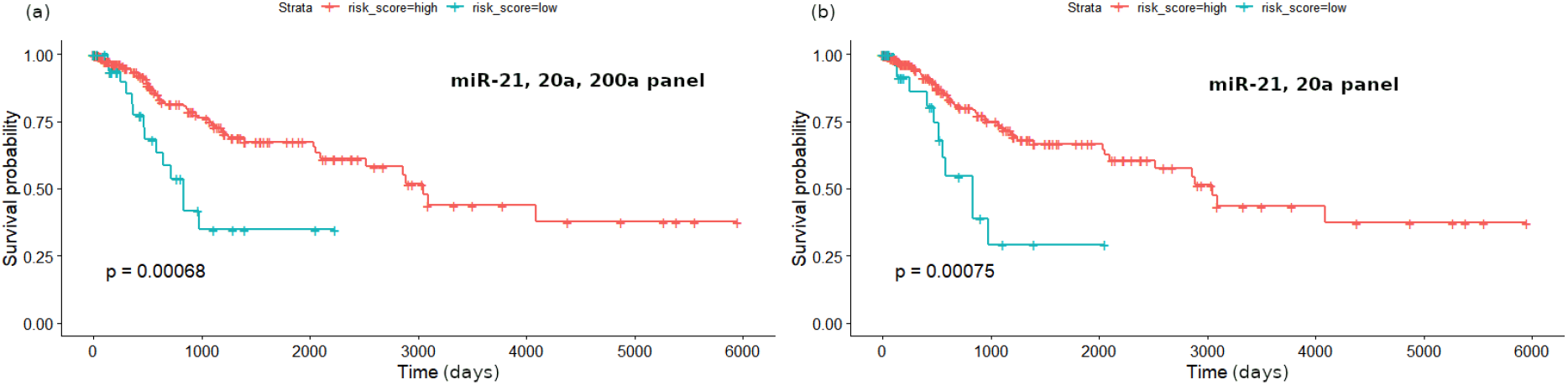
(a) Kaplan-Meier curves of panel with the three chosen biomarkers: hsa-miR-20a-5p, hsa-miR-21-5p and hsa-miR-200a-3p (b) Kaplan-Meier curves of panel with the two chosen biomarkers: hsa-miR-20a-5p and hsa-miR-21-5p.

### 3.3 Toehold Switch Design

Two toehold switches were designed to target the identified miRNA biomarkers, hsa-miR-21-5p and hsa-miR-20a-5p (provide supplementary info on miRNA sequences, toehold sequences, and corresponding DNA sequences). The folding of the transcribed RNA of the designed sequences was predicted using NUPACK [42] and ViennaRNA [28], and the secondary structure was consistent with the toehold conformation (Fig. 5a, b). Neither of the two structures have any rogue base-pairings in the toehold domain. In addition to the secondary structure, the minimum free energy and the dynamic range of the designed toehold switches are key parameters in the effectiveness of the system. The minimum free energies of the hsa-miR-21-5p toehold switch and the hsa-miR-20a-5p toehold switch were estimated to be −21.20 kcal/mol and −19.20 kcal/mol respectively, suggesting a stable toehold conformation, ready for interaction with the cognate miR-antimiR complex.

**Fig 5:**
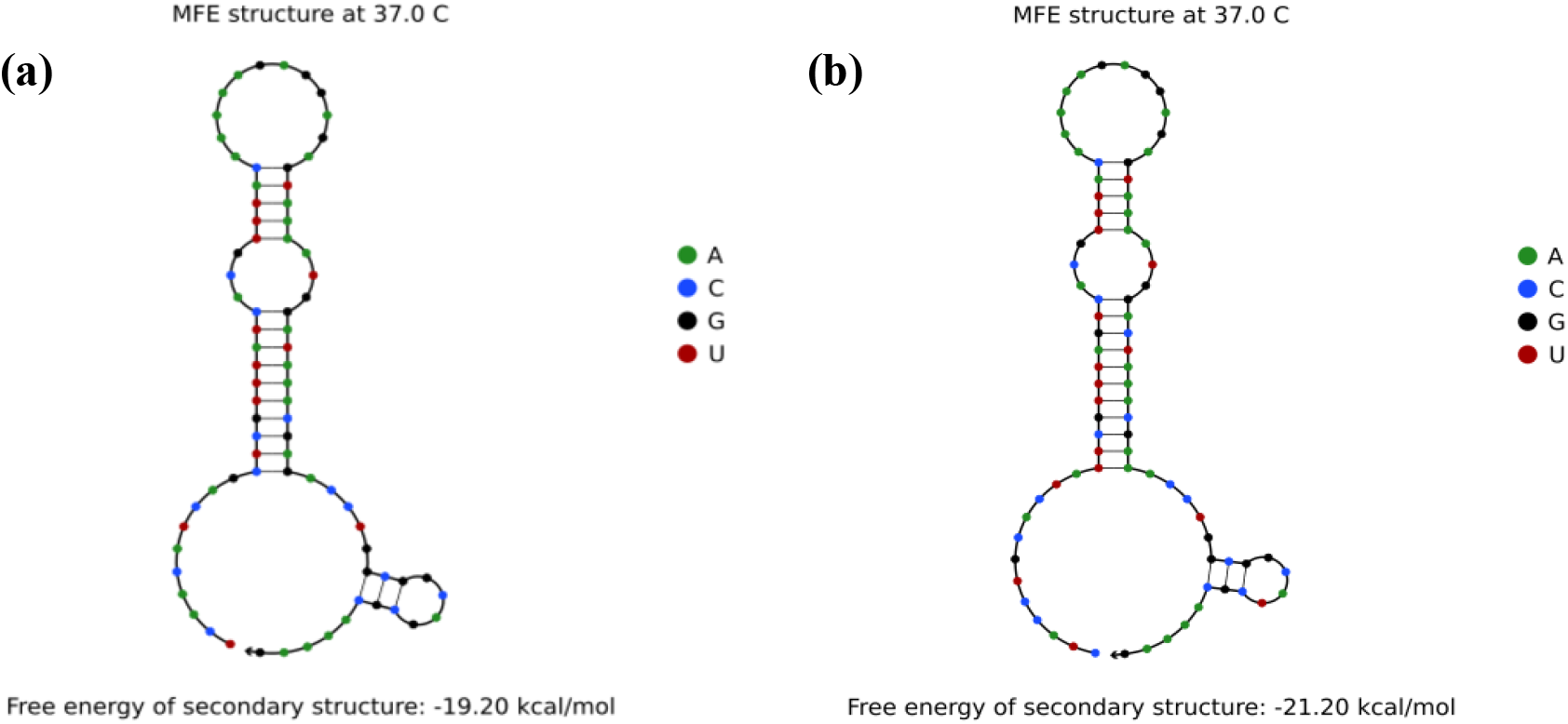
(a) MFE structure of Second Generation Toehold Switch for hsa-miR-21-5p, (b) MFE structure of Second Generation Toehold Switch for hsa-miR-20a-5p shown in NUPACK.

Feature selection for our machine learning model yielded six non-redundant features, all thermodynamic in origin:

1. Overall MFE of the toehold switch construct
2. Frequency of the MFE structure in the ensemble
3. Net MFE (difference in energy between the MFE of the switch-trigger complex and the sum of individual MFE values of the switch and trigger)
4. MFE of the bottom region containing the toehold domain, bottom stem and the horizontal linker
5. MFE of the RBS-linker domain, starting from the primary loop till the end of the construct
6. Specific heat of the toehold switch construct

It is notable that net MFE which is an energetic feature arising from the interaction of the second-generation toehold with the miRNA-antimiR hybrid has emerged as a key feature in the modelling process. The significance of the coefficients of the resulting multivariate linear regression model are noted in Table 6. The model yielded an adj. R^2^ ~ 0.59 on the test set. With this validation, we applied the model to the designed toehold switches for the two identified miRNA biomarkers. The toehold switch for hsa-miR-21-5p yielded a predicted dynamic range of 52.4, while that of hsa-miR-20a-5p yielded 100.8, both values indicating potential robust performance of the designed biosensors. The model is freely available for use in toehold-based synthetic biology [29]. All the designed toehold switch sequences and genetic circuits are available in File S4, and the BioBricks submissions are noted in File S5.

**Table 6.** Significance of the selected features in the constructed multivariate linear regression model (All features were significant)

| S. No. | Feature | Coefficient | p-value |
| --- | --- | --- | --- |
| 1 | Overall MFE | 16.39 | 1.7E-03 |
| 2 | Bottom_region_MFE | -32.81 | 1.8E-11 |
| 3 | RBS_linker_MFE | 19.21 | 2.8E-06 |
| 4 | Net_MFE | 5.24 | 3.3E-03 |
| 5 | Freq_MFE_struct (frequency) | -327.33 | 2.4E-02 |
| 6 | Overall_SH_37 (specific heat) | 68.21 | 6.3E-05 |

### 3.4 Reaction Modelling

We employed reaction network mass action kinetics to computationally evaluate the performance of the designed toehold switches as biosensors of miRNA biomarkers (available as executable SBML model in github repository; submitted to EBI BioModels Repository). Before understanding the reaction kinetics of our designed second generation toehold switches, we initially decided to emulate the model proposed by iGEM CLSB-UK (2017) [40] to obtain an insight of their kinetic model on first generation toehold switches and anticipate the plausible errors that could be encountered during our simulations (File S6)

#### 3.4.1 Generalization to Second generation Toehold Switches

We performed NUPACK runs to obtain the structures of hsa-miR-21-5p and anti-hsa-miR-21-5p [42], and analyzed the equilibrium of their interaction at 18 °C. Using the equilibrium concentrations of the complex, hsa-miR-21-5p, and anti-hsa-miR-21-5p (99.97 nM, 0.0264 nM, and 0.0264 nM respectively), the k_miR_antimiR_f_ was estimated to be in the order of 10^5^ nM^−1^. The concentration profiles for hsa-miR-21-5p, anti-hsa-miR-21-5p and their complex are shown in Fig. 6(a).

**Fig 6.**
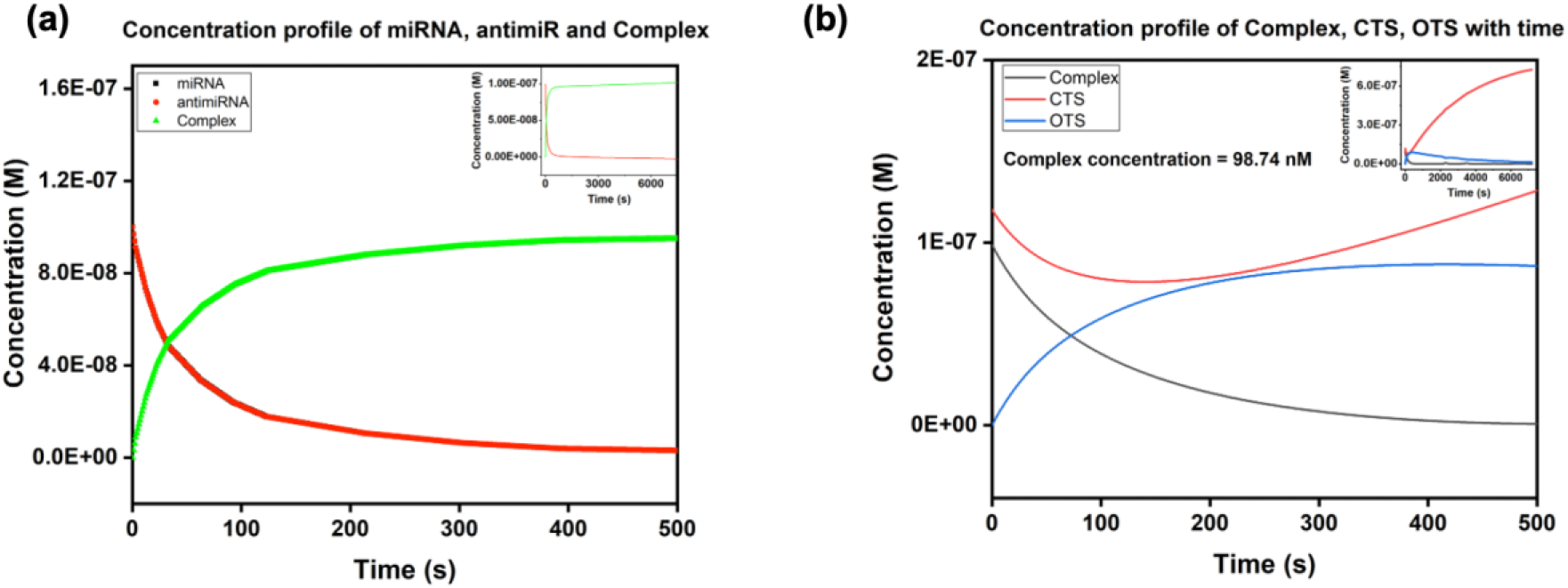
(a) Concentration profiles of miR, antimiR and the complex. It is seen that miR and antimiR profiles are identical. (b) Concentration profiles of the complex, CTS and OTS in a 2 h time frame with a miR-antimiR binding rate constant of 10^5^ mol^−1^s^−1^. The first 500s are highlighted with an inset over the simulation time of 7200s.

The simulation time was set to 7200 s (2 h) allowing for miRNA-antimiR complex formation. From the profile in Fig. 6(a), it is clear that the complex formation occurs rather rapidly and both the complex and the miR/anti_miR attain saturation by 500s. We used an incubation time of 8 minutes for the cell-free transcription of the genetic construct to yield significant quantities of the RNA toehold switch [41].

The intensity of GFP fluorescence is the readout from the proposed biosensor. A GFP maturation reaction was included in the model with a maturation factor ~ 0.2/min. To obtain the raedout in terms of fluorescence units, we utilized the scaling factor of 79.429 proposed by iGEM Valencia_UPV [49]. The initial DNA concentration was set to 223.5 nM, following established experimental protocols.

From the concentration profile seen in Fig. 6(b), the CTS concentration was initially ~118 nM due to the transcriptional incubation time. The complex concentration was found to be 98.74 nM upon solving the differential equations. The CTS and complex rapidly hybridize to open the CTS, evident from the significant drop in their concentrations along with a rise in the OTS concentration. At about 500s, the complex gets completely consumed, following which the OTS begins to decay, while CTS continues to rise due to unhindered transcription. Upon looking into the dynamics of GFP and mature GFP during the initial 500s, translation of GFP occurs with increase in OTS concentration followed by a lazy maturation of the GFP.

The final phase of our modelling included the process of obtaining trends for fluorescence intensity with variations in complex concentration - from 100 pM to 100 nM. We plotted the intensity at 2 h interval against the complex concentration and obtained a linear fit between 10 nM to 100 nM with a goodness-of-fit R^2^ = 0.9993. This showed that the model is sensitive to changes in the concentration of the trigger miRNA biomarker, satisfying a necessary biosensor property [50–52]. We repeated the process for a time interval of 4h where a similar trend was obtained (R^2^ = 0.9992) (Fig.7).

**Fig 7.**
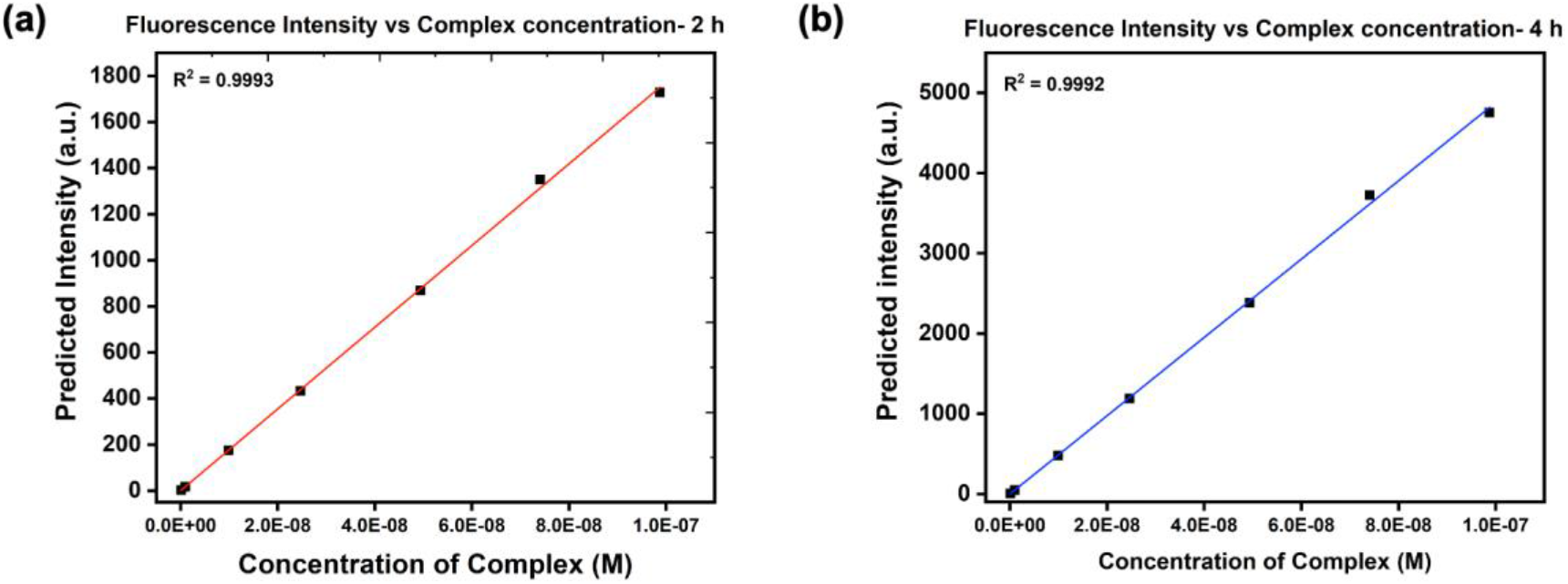
The fluorescence intensity varies linearly with complex for a concentrations between 10 nM and 100 nM over (a) 2h and (b) 4h.

Table 7 provides the concentration trajectories of various species and the predicted fluorescence intensity readout for different miRNA-antimiR complex concentrations assuming a 2-hr reaction time. The intensity values for an extended 4-hr reaction time are provided in Supplementary Section S2.

**Table 7.**
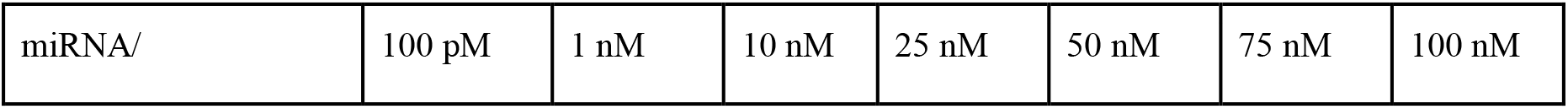
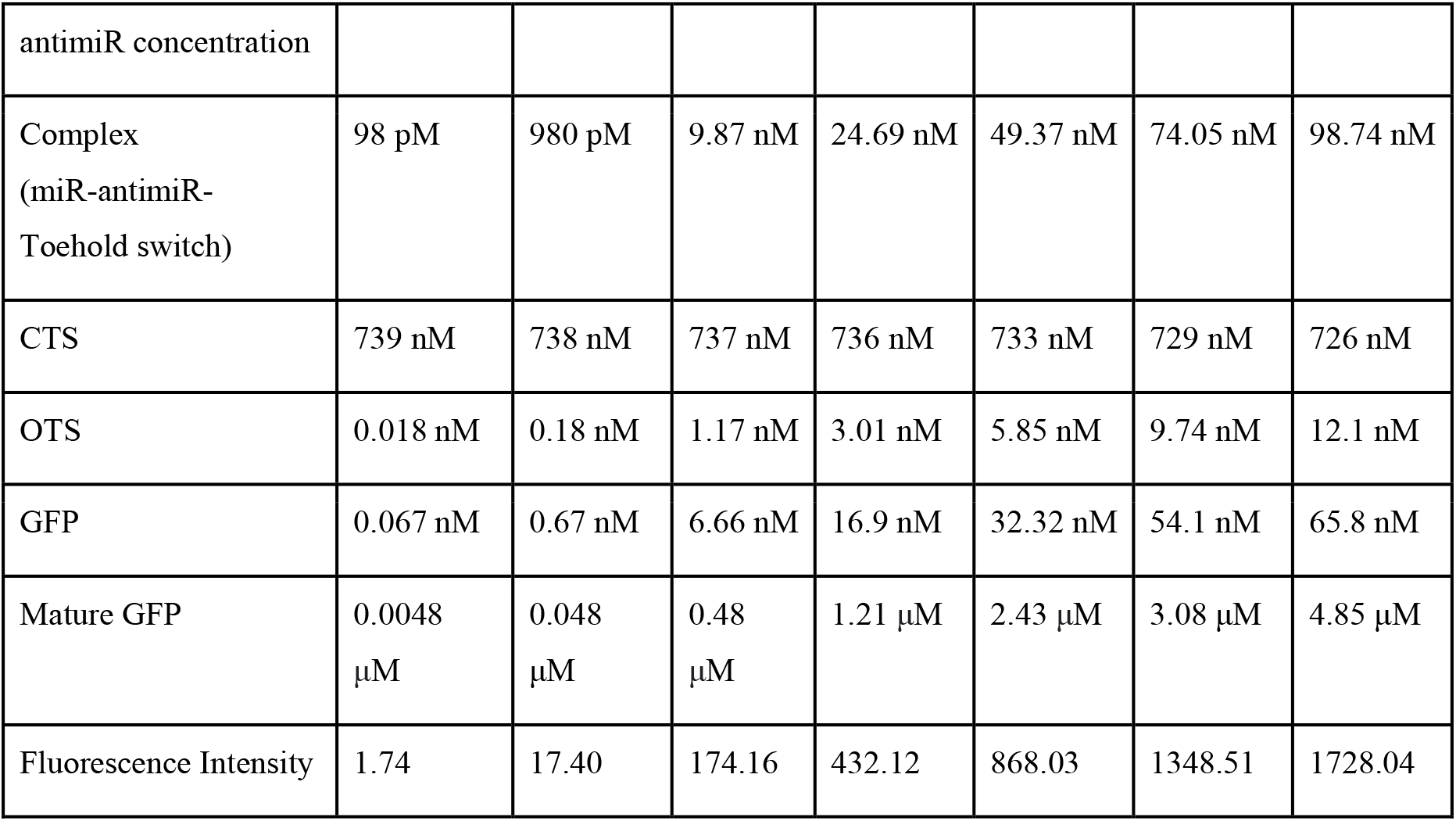
Simulated fluorescence intensity variations with different concentrations of the complex over a 2h period.

|  |  |  |  |  |  |  |  |
| --- | --- | --- | --- | --- | --- | --- | --- |
| miRNA/ | 100 pM | 1 nM | 10 nM | 25 nM | 50 nM | 75 nM | 100 nM |
| antimiR concentration |  |  |  |  |  |  |  |
| Complex<br>(miR-antimiR-Toehold switch) | 98 pM | 980 pM | 9.87 nM | 24.69 nM | 49.37 nM | 74.05 nM | 98.74 nM |
| CTS | 739 nM | 738 nM | 737 nM | 736 nM | 733 nM | 729 nM | 726 nM |
| OTS | 0.018 nM | 0.18 nM | 1.17 nM | 3.01 nM | 5.85 nM | 9.74 nM | 12.1 nM |
| GFP | 0.067 nM | 0.67 nM | 6.66 nM | 16.9 nM | 32.32 nM | 54.1 nM | 65.8 nM |
| Mature GFP | 0.0048<br>μM | 0.048<br>μM | 0.48<br>μM | 1.21 μM | 2.43 μM | 3.08 μM | 4.85 μM |
| Fluorescence Intensity | 1.74 | 17.40 | 174.16 | 432.12 | 868.03 | 1348.51 | 1728.04 |

## 4. Discussion

Geraldi *et al.* highlighted the importance of synthetic biology in developing portable *in vitro* based diagnostic kits [53]. Toeholds embedded in genetic circuits enable the measurement of the intensity of expression of a downstream reporter protein – for e.g, GFP – that could yield a precise quantification of the trigger itself. Pardee has reviewed the potential of freeze-dried cell-free (FD-CF) systems in health care for diagnostic and sensing purposes, given their biosafe mode [54], and Tinafar *et al* reviewed cell free systems and the promising opportunity for the rational design and manipulation of biological systems in relation to cell based systems [55]. Fabricating microfluidic devices with our designed genetic circuits in a cell-free system followed by rigorous testing and validation would yield a point-of-care portable diagnostic tool for the real-time detection of early-stage cervical cancer.

Towards this end, uniting theory and data is key, and Figure 8 encapsulates the methodology we have adopted here, in two parts. The end-to-end computational approach that we have employed in this study is illustrated in Fig. 8(a). It broadly includes four phases, namely identification of biomarker signatures, design of the biosensory element - toehold switch followed by the synthetic genetic circuit design comprising modular constructs and finally validating a proof-of-concept of the designed synthetic circuit using predictive modelling strategies. This approach could be generalised to the design of portable, highly sensitive and specific biosensors for any condition and especially infectious diseases. The current nCoV-SARS2 pandemic has highlighted the need for constant vigil for outbreak of infections. Such emerging agents require accurate, rapid, and scalable detection platforms, all requirements for which toehold biosensors are well-suited. Fig. 8(b) illustrates a possible scheme for the development and production of such a toehold biosensor. Here the biomarker would be an optimal genomic fragment signature, unique to the pathogen but not in the host genome. The toehold construct serves to signal the presence of the infectious agent that would then release the expression of the reporter gene, yielding a simple visual readout in real-time within minutes. The proposed technique upon execution would generate a simple, reliable, rapid, portable and affordable printed diagnostic tool that could be made available at a global level in a point-of-care setting outside clinical diagnostic laboratory conditions. This simple biosafe product would be an asset in the field, especially in remote locations where it could help in curtailing the transmission and limiting the reproductive ratio of the infectious agent. Such a product would be vital in low-resource settings since it is sterile, abiotic and active for a year at room temperature. Moreover the product is easily scalable, saving the lives of countless frontline corona warriors and resulting in improved patient outcomes. Extension of more than two biomarkers would allow for the simultaneous detection of, say, a family of coronaviruses, an imperative in our times. Thus, end-to-end science is expected to overcome the limitations of ad-hoc diagnostic devices, thereby enabling affordable diagnosis in pandemic situations and other health emergencies.

**Fig 8.**
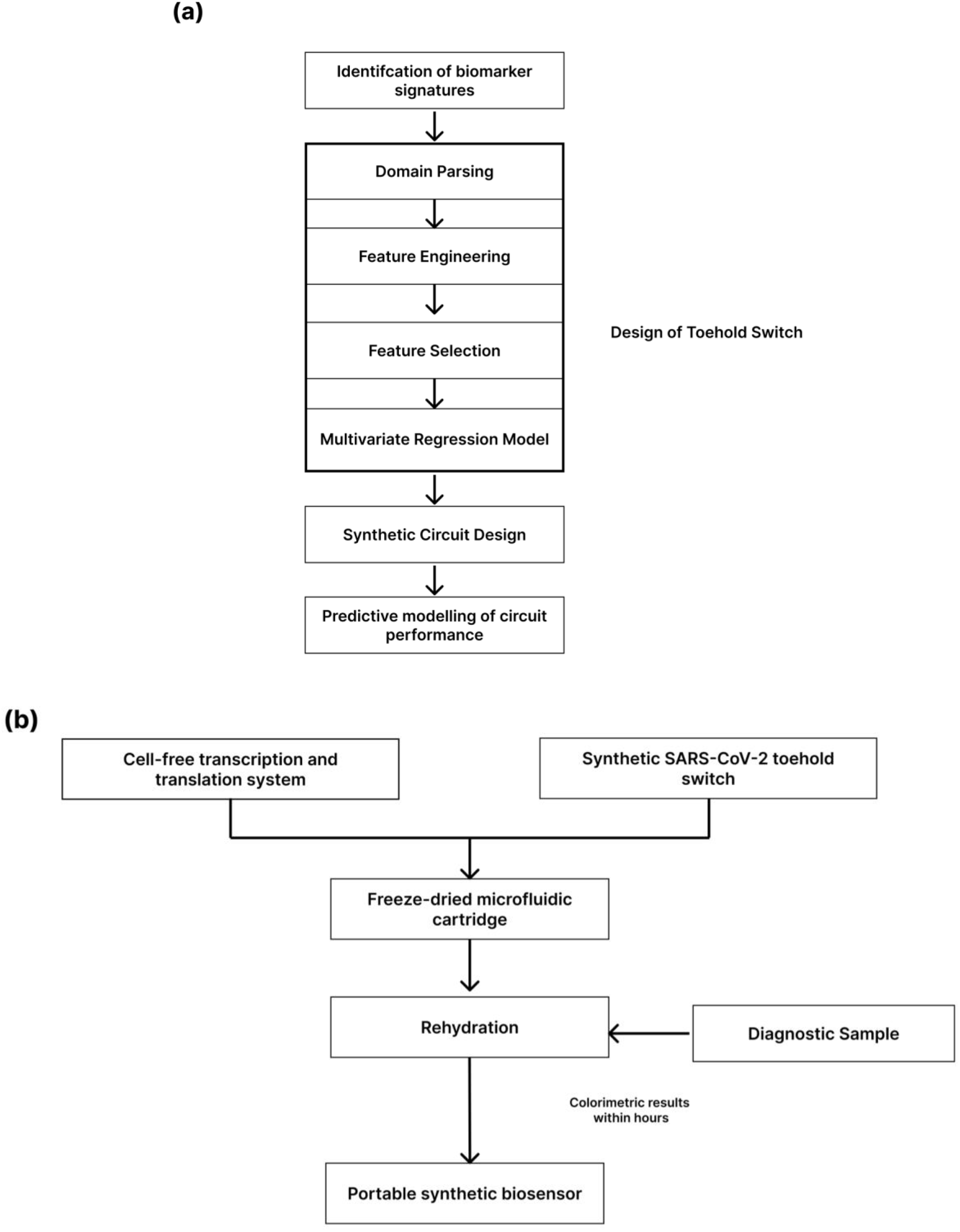
(a) General scheme representing the end-to-end computational approach for design of toehold switches based biosensors (b) Scheme representing the modus operandus for designing and developing biosensors working on a riboregulator based sensing platform where toehold switches and SARS-CoV-2 viral genomic content act as the biosensory element and analyte, respectively.

## 5. Conclusion

Cervical cancer is a major public health issue, but an addressable one with more effective diagnosis technologies. Early detection results in better treatment and greater compliance. In this work, three DBTL iterations were used to generate a workflow for *in silico* biomarker detection, sensor design and systems dynamics modelling. In the first cycle, we identified miRNAs that were significantly differentially expressed for early-stage cervical cancer using TCGA data, and identified the optimal two miRNA biomarkers. In the second cycle, we designed second generation toehold switches and antimiR for the two miRNAs. In this course, we developed a machine learning model for toehold dynamic ranges that provided an adj.R^2^ = 0.59, a major improvement in efforts in this direction. Finally we simulated the reactions in the cell free system to study their kinetics. From our observations, fluorescence intensity followed a linear range between 10 nM and 100 nM after which we observed a saturation. End-to-end automation of the computational ideas in our workflow would essentially democratize detection of emerging infectious diseases and life-threatening conditions with a readily transportable instrument based on a real-time visual readout.

## Author contributions

Conceived and designed the work: AP, SS, PRS. Performed research: PRS, SN, SS, SM, RRS, TS. Analyzed the results: PRS, SS, SN, SM, RRS, AP. Writing- first draft: PRS, SS, SN, SM, RRS. Writing- editing and revision: AP. Supervision & project administration: AP.

## Acknowledgements

We are grateful to SASTRA Deemed University for financial and infrastructural support to SASTRA iGEM 2019 Team (https://2019.igem.org/Team:SASTRA_Thanjavur). A.P. would like to acknowledge support from DST-SERB EMR/2017/000470.

